# Temporal Dynamics of Transcriptional Responses to Repeated mRNA Vaccination: Insights from Third Dose Profiling

**DOI:** 10.1101/2025.01.09.631990

**Authors:** Darawan Rinchai, Sara Deola, Basirudeen Syed Ahamed Kabeer, Giusy Gentilcore, Mohammed Toufiq, Lisa Mathew, Li Liu, Fazulur Rehaman Vempalli, Ghada Mubarak, Stephan Lorenz, Tracy Augustine, Nicholas Van Panhuys, Davide Bedognetti, Jean-Charles Grivel, Damien Chaussabel

## Abstract

mRNA vaccines have played a crucial role in combating the COVID-19 pandemic, but the long-term dynamics of immune responses to repeated vaccination remain poorly understood. In this study, we extend our previous work on first and second dose responses by characterizing the immune signatures elicited by a third dose of COVID-19 mRNA vaccines using high-resolution temporal profiling of blood transcriptomes collected daily for 9 days post-vaccination. We observed distinct patterns of gene expression related to interferon responses, inflammation, erythroid cell signatures, and plasmablast activity across the three doses. While the first dose elicited a modest response primarily characterized by interferon signaling, the second dose induced a robust, polyfunctional response. The third dose, administered approximately nine months later, maintained this polyfunctional character and matched the second dose in magnitude, though with distinct temporal dynamics. The interferon component peaked on day 2 (similar to the first dose) rather than day 1 (as seen in the second dose), while the erythroid signature showed a markedly different trajectory, with sustained elevation rather than decrease over the following week. Notably, we observed a progressive amplification of the plasmablast response across the three doses, with an earlier peak (day 4) compared to other vaccines, potentially a unique feature of mRNA vaccines. These findings demonstrate that the heightened, polyfunctional responsiveness induced by the second dose is robustly maintained even after a prolonged interval, suggesting effective immune memory. Our results contribute to understanding mRNA vaccine-induced immunity, with implications for optimizing booster strategies and developing next-generation vaccines.

## INTRODUCTION

mRNA vaccines, a new and emerging vaccine platform, have been approved for large-scale use in humans at the end of 2020, primarily in response to the SARS-CoV-2 pandemic (1,2). Initial trials have demonstrated their superior protection levels against SARS-CoV-2 infection compared to more traditional vaccine approaches. However, their effectiveness in practice has been somewhat reduced due to the emergence and rapid spread of highly contagious variants (3,4). Despite this, mRNA vaccines have been instrumental in significantly reducing the risk of developing severe symptoms and effectively curbing mortality (5,6). As this technology continues to evolve, the range of applications for mRNA vaccines is expanding rapidly, encompassing other infectious diseases and cancer. However, the characteristics of the immune response elicited by this novel vaccine platform and how it relates to its efficacy are not yet entirely defined.

To date, the immunological profile of mRNA vaccine responses has been characterized through various approaches, including omics technologies and traditional immunological assays. Studies have shown that mRNA vaccines induce strong humoral and cellular immune responses, with high levels of neutralizing antibodies and robust T cell responses (7). Notably, mRNA vaccines have been found to elicit a broader range of memory B cells compared to natural infection, potentially conferring enhanced protection against viral variants (8). Transcriptomic studies have revealed early activation of innate immune pathways, including interferon and inflammatory responses, which may contribute to the vaccines’ efficacy (9). Additionally, single-cell RNA sequencing has identified specific immune cell subsets, such as cross-reactive T cells, that are preferentially expanded following mRNA vaccination (10,11). Despite these insights, the long-term dynamics of the immune response to repeated mRNA vaccination and the mechanisms underlying their exceptional efficacy remain areas of active investigation.

In our previous work, we employed bulk blood transcriptomics at a high temporal frequency to explore the immune response to COVID-19 mRNA vaccines (12). We reported a striking enhancement of day-1 responses post-administration of the second vaccine dose compared to the first. This early response was characterized as polyfunctional, dominated by innate signatures such as interferon, inflammation, and erythroid cells. By day 5, we observed an adaptive signature marked by plasma cell signatures, which correlated with the serological response measured at days 14 and 28 post-vaccination. In the present study, we applied the same high-frequency blood transcriptomics approach to characterize the response to the third vaccine dose. This allowed us to investigate whether the immediate response observed post-second dose would remain amplified months later. Moreover, by analyzing the transcriptional profiles across all three doses, we aimed to uncover the evolving dynamics of the immune response to repeated mRNA vaccination. This comprehensive analysis not only provides insights into the long-term efficacy of mRNA vaccines but also contributes to our understanding of how the immune system adapts to sequential exposures to the same antigen. Our findings have implications for optimizing booster strategies and may inform the development of next-generation mRNA vaccines for various applications beyond COVID-19.

## RESULTS

### Profiling blood transcriptome trajectories and serological responses post-third COVID-19 mRNA dose

To gain a comprehensive understanding of the immune response induced by repeated administration of COVID-19 mRNA vaccines, we sought to obtain high temporal resolution blood transcriptome profiles from subjects receiving a third dose. This approach builds upon our previous work characterizing responses to the first and second doses, allowing for a more complete picture of the evolving immune response.

We generated RNA-sequencing data from samples collected at high temporal frequencies from 17 subjects who received a third dose of COVID-19 mRNA vaccine (**Figure 1**). This cohort included 10 subjects from our earlier study on first and second dose responses, and 7 newly enrolled subjects. The sampling schedule mirrored our previous protocol (see **Figure 1** and Methods), with blood samples collected immediately before vaccination, on the day of vaccination, and daily for 9 consecutive days post-vaccination. Additional samples were collected on days 14 and 28 for serological profiling. Our cohort’s COVID-19 vaccination and infection history is detailed in **Supplementary File 1**. Notably, five subjects were infected prior to their first dose, three between their second and third doses, and five after their third dose. Two subjects who developed symptoms shortly after vaccination were excluded from the analysis. We employed a self-collection protocol allowing participants to collect small blood volumes (50 µl) at the required frequency (described in Methods and (13)). Blood transcriptome profiles were generated using a cost-effective 3′-biased RNA-sequencing protocol optimized for low RNA input. Antibody profiles were produced using capillary blood specimens collected via Volumetric Absorptive Micro Sampling and analyzed with a custom multiplexed bead array (see Methods for details).

**Figure 1:**
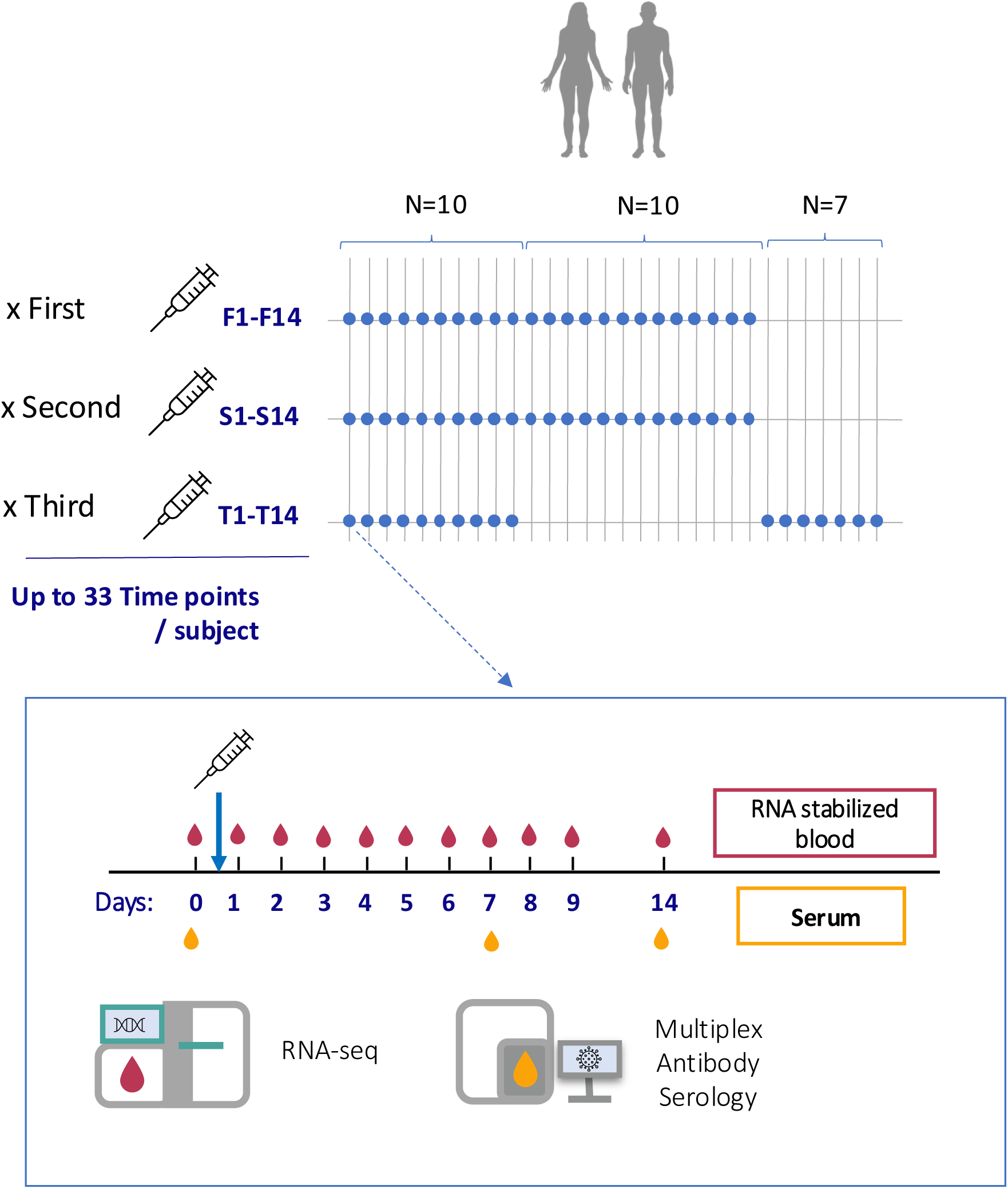
Study design and sampling schedule for profiling immune responses to COVID-19 mRNA vaccine doses. This figure illustrates the study design and sampling schedule for profiling immune responses to three doses of COVID-19 mRNA vaccines. (A) Cohort composition and sampling timeline. The study included 30 adult subjects in total: 10 subjects (N=10) who received all three doses and were sampled throughout, 13 subjects (N=13) who were sampled for the first and second doses only, and 7 newly recruited subjects (N=7) who were sampled for the third dose only. Blue dots represent sampling time points (F: First dose, S: Second dose, T: Third dose). (B) Detailed sampling schedule for each vaccine dose. Blood samples were collected immediately before vaccination (day 0), on the day of vaccination, and daily for 9 consecutive days post-vaccination. Additional samples were collected on days 14 and 28. Red droplets indicate RNA-stabilized blood samples collected for transcriptome analysis (RNA-seq). Orange droplets indicate serum samples collected for antibody profiling. (C) Sample processing workflow. RNA-stabilized blood samples were used for RNA-seq analysis to generate transcriptome profiles. Serum samples were analyzed using a multiplex antibody serology assay to measure vaccine-induced antibody responses. This high-frequency sampling approach allows for detailed characterization of the kinetics of immune responses following each dose of the mRNA vaccine, with up to 33 time points per subject across the three doses.

RNA-seq profiling data have been deposited in GEO (GSE281864) for future reuse. This effort yielded high temporal resolution blood transcriptome data post-third dose for 17 individuals, with 10 of these having data available across all three vaccination rounds (36 time points in total). UMAP plotting of transcriptomic profiles revealed that responses generated largely segregated based on whether the subjects were receiving the first, second or third dose of vaccine (**Figure 2A**). The antibody profiles we examined showed a reaction to the stable trimer of the Spike protein, its receptor-binding domain, the Nucleo and Envelope proteins of SARS-CoV2. We dissected the reactivity to each antigen, measuring the total IgG, total IgA, and IgM, as well as more specific IgG and IgA subtypes. It was observed from the antibody profiling that there was an increase in antibody levels in the plasma of subjects post-third vaccine dose (**Figure 2B and C**), including antibodies specifically targeting the SARS-CoV-2 Spike protein, which are targeted by the COVID-19 mRNA vaccine. We found no reactions to the Envelope protein, although we did observe some cross-reactivity with the SARS Spike protein (**Figure 2C**). As anticipated, higher levels of antibodies were elicited after the first vaccine dose in subjects who had prior infection with the virus. These observations align with past studies that have characterized the antibody response to COVID-19 mRNA vaccines (14).

**Figure 2:**
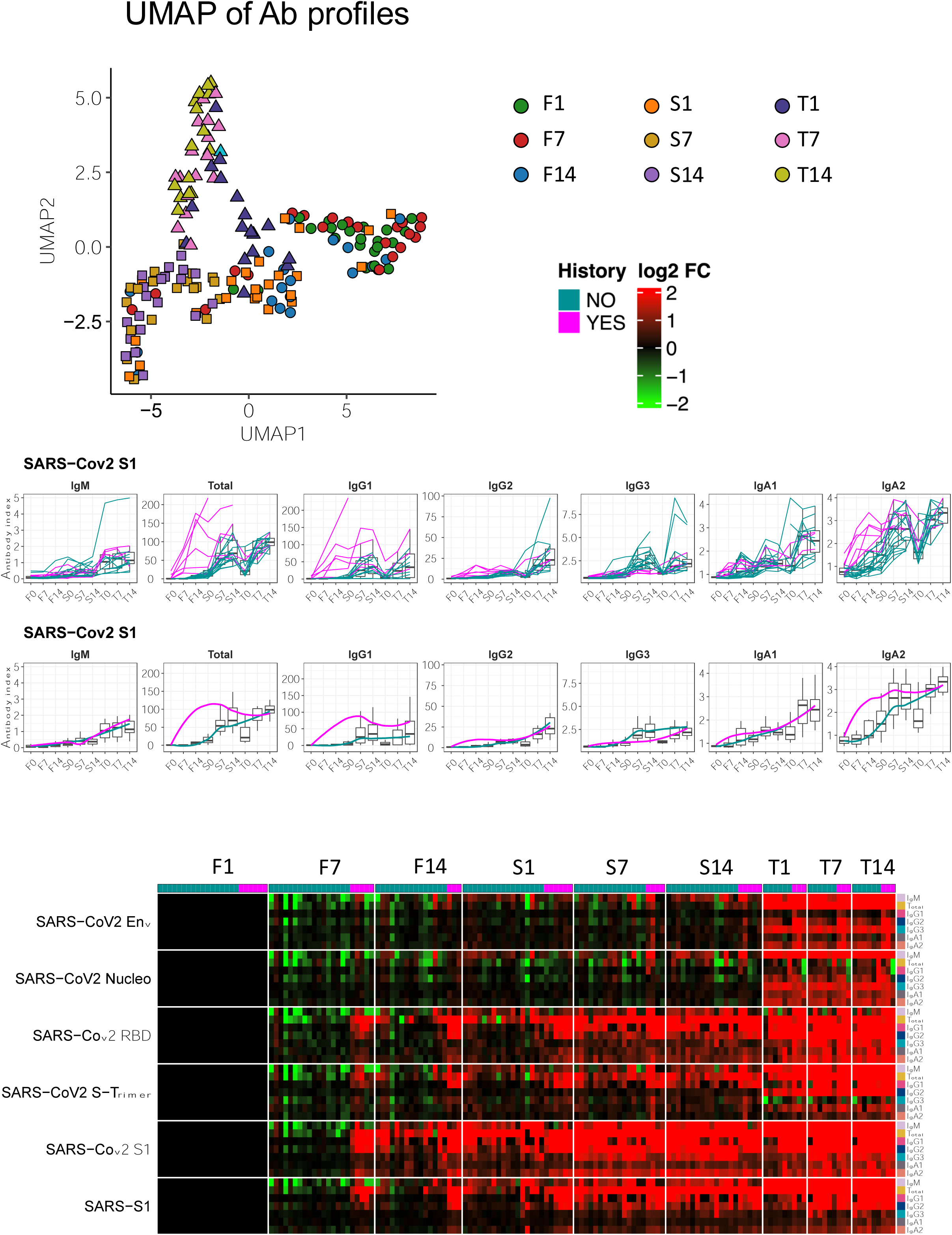
Antibody profiling and transcriptional changes across three doses of COVID-19 mRNA vaccine. A) UMAP plot of transcriptome profiles. Each point represents a sample, colored by time point (C1D01-C3D14) and shaped by dose (circles for first dose, triangles for second dose, squares for third dose). B) Antibody profiles for SARS-CoV2 S1 antigen. Top panel shows individual subject responses, bottom panel shows averaged responses. Each line graph represents a different antibody isotype (IgM, Total, IgG1, IgG2, IgG3, IgA1, IgA2). X-axis shows time points (F0-F14: first dose, S0-S14: second dose, T0-T14: third dose). Y-axis represents antibody index. Pink lines in the top panel indicate subjects with prior COVID-19 infection history. C) Heatmap of antibody responses to various SARS-CoV2 antigens. The color gradient in the legend represents log2 fold change, with pink indicating prior COVID-19 infection history. Rows represent different antigens (Env, Nucleo, RBD, S-Trimer, S1) and SARS-S1 across various isotypes. Columns represent time points across three doses (C1-C3) and three time points (D01 = pre-vaccination, D07 = 7 days post-vaccination, D14 – 14 days post-vaccination). Color intensity indicates the strength of antibody response, with red showing stronger responses and green showing weaker responses, relative to the pre first dose baseline (C1D01). Each small square within a time point represents an individual subject.

This comprehensive dataset provides an unprecedented opportunity to examine the evolution of immune responses across multiple doses of mRNA vaccines, potentially revealing insights into the mechanisms of vaccine-induced immunity and the effects of repeated vaccination.

### The third COVID-19 mRNA Vaccine Dose Elicits a Potent Polyfunctional Response, Mirroring that Seen Post-Second Dose

To understand how the immune response evolves with repeated mRNA vaccination, the response post-third dose was benchmarked against those observed post-first and second doses. Blood transcriptional profiles were analyzed at the module level using the BloodGen3 modular repertoire and analysis tools, as previously employed for the first and second dose response analysis (see Methods for details and (12,15)).

Of the 382 modules in this repertoire, 266 responded on day 1 post-third dose (defined as at least 15% of constituent transcripts showing gene expression differences). Among these, 98 modules showed increased transcript abundance, while 168 exhibited decreased abundance (**Figure 3A**). This response closely paralleled that observed post-second dose (261 responsive modules, 91 increased, 170 decreased) but significantly exceeded the response post-first dose (7 responsive modules). The third-dose response appeared more robust than the second dose on days 2 and 3 post-vaccination, with 165 and 42 responsive modules, respectively.

**Figure 3:**
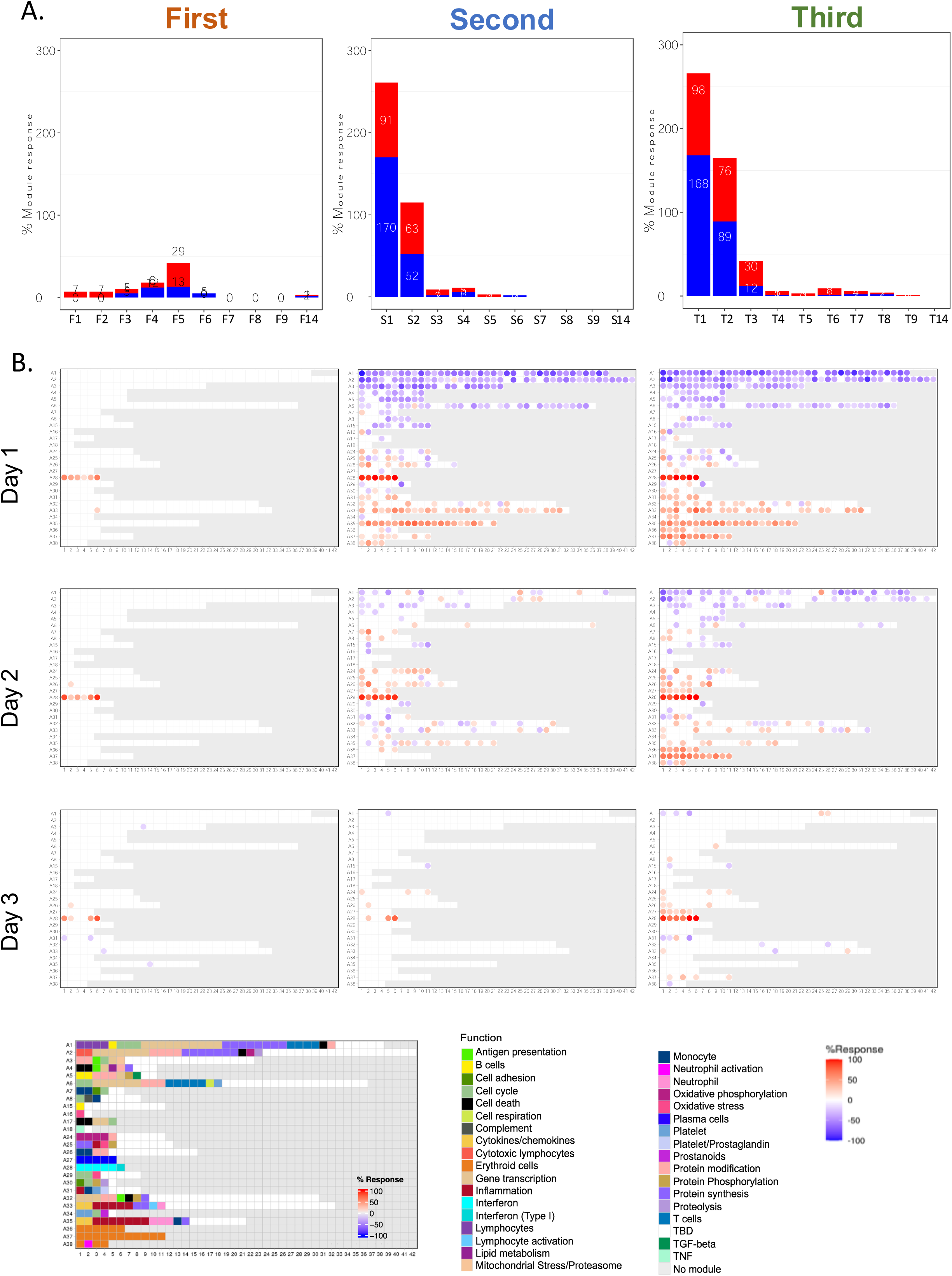
Comparative analysis of transcriptional responses to first, second, and third doses of COVID-19 mRNA vaccine. (A) Bar graphs showing the number of responsive BloodGen3 modules at each time point following administration of the first, second, and third vaccine doses. Red bars indicate modules with increased transcript abundance, while blue bars represent modules with decreased abundance. The x-axis shows days post-vaccination, and the y-axis represents the number of responsive modules. (B) Fingerprint grid plots illustrating module-level responses on days 1, 2, and 3 post-vaccination for each dose. Each spot on the grid represents a module, with its position fixed across all plots. Red spots indicate increased transcript abundance, blue spots indicate decreased abundance, and the intensity of color represents the magnitude of change. Modules are arranged in rows corresponding to module aggregates (A1 to A38). The bottom panel provides a color-coded functional annotation key for the modules.

Qualitative differences were highlighted using fingerprint grid maps designed for visualizing the BloodGen3 module repertoire (**Figure 3B**). The first dose elicited an interferon response signature, evidenced by a six-module increase in aggregate A28. This transcript abundance also increased post-second and third doses, with notable changes for A28 modules continuing through day 3 post-third dose. On a broader scale, the responses to the second and third doses appeared similar. Decreased abundance was observed in module aggregates typically associated with lymphocytic cell populations (A1 through A6), while increased transcript abundance was noted in modules linked with inflammation (A33, A35) and erythroid cells (A36, A37, A38). This increase was more pronounced post-third dose than post-second dose, particularly on day 2 post-vaccination. A subtle distinction was the A27 signature, associated with plasmablast responses, noticeable on days 2 and 3 post-third dose, but not post-second dose.

These findings indicate that the post-third dose response strongly aligns with the potent and polyfunctional response observed post-second dose, marking a clear departure from the modest response post-first dose. This suggests a potential cumulative effect of repeated mRNA vaccination on the immune response, though the partial subject overlap between second and third dose cohorts necessitates cautious interpretation.

### The interferon response to the third vaccine dose exhibits kinetic and amplitude variations compared to responses post-first and second doses

Interferon responses are critical components of the innate immune response to vaccines and are known to influence the development of adaptive immunity. Understanding how these responses evolve across multiple vaccine doses can provide insights into the mechanisms of vaccine-induced protection.

We analyzed changes in the interferon signature across all collection time points following the first, second, and third doses of the COVID-19 mRNA vaccine. We focused on the A28 module aggregate, which comprises six interferon-related modules previously identified in our BloodGen3 repertoire.

The kinetics and amplitude of the interferon response varied markedly across the three vaccine doses (**Figure 4A**). After the first dose, the response peaked on day 2, while after the second dose, it showed a sharp peak on day 1. Interestingly, the response to the third dose resembled that of the first dose in timing, peaking on day 2, but matched the second dose in amplitude. Further analysis revealed two distinct components of the interferon signature: S1 (modules M8.3, M10.1, and M15.127) and S2 (modules M13.17, M15.64, M15.86) (**Figure 4B-C**). These components exhibited different patterns of response across the three doses. After the first and third doses, S1 modules peaked on day 2, while S2 modules peaked on day 1. In contrast, after the second dose, both S1 and S2 peaked simultaneously on day 1.

**Figure 4:**
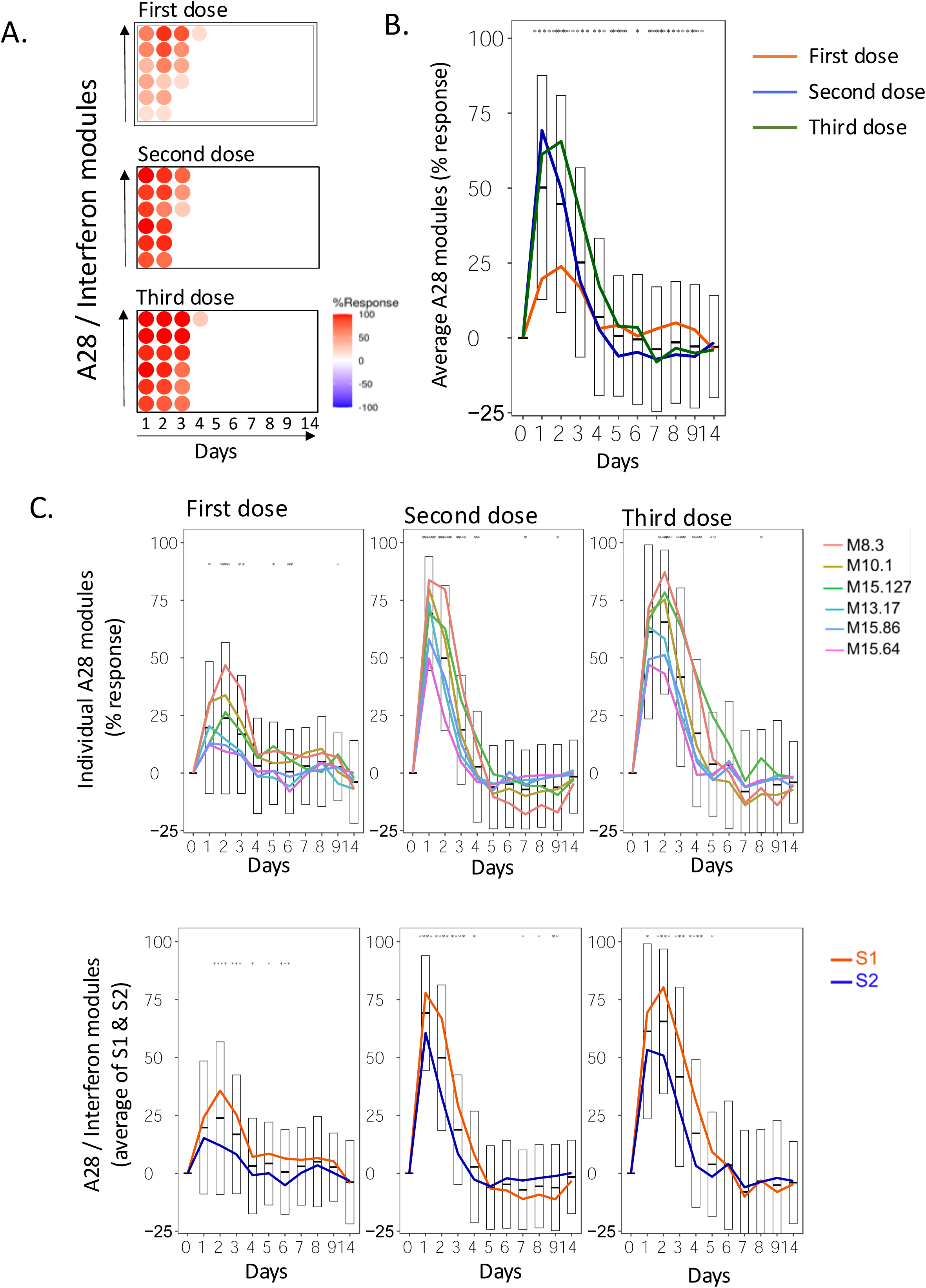
Interferon response patterns across three doses of COVID-19 mRNA vaccine. (A) Heatmaps showing the response of A28 interferon modules over time for each vaccine dose. Each row represents one of the six A28 modules, and columns represent days post-vaccination. Color intensity indicates the percentage of responsive genes within each module, with red indicating increased expression and blue indicating decreased expression. (B) Line graph showing the average response of all A28 modules over time for each vaccine dose. Lines represent the mean response, with error bars indicating the standard deviation. Asterisks above the graph denote statistically significant differences between doses at each time point, with * p<0.05, ** p <0.01, *** p < 0.001. (C) Line graphs showing individual A28 module responses (top row) and the average responses of S1 and S2 module subsets (bottom row) for each vaccine dose. Each colored line represents a different module or module subset. The gray bars in the background represent the range of responses across all modules. Asterisks above the graph denote statistically significant differences between doses at each time point, with * p<0.05, ** p <0.01, *** p < 0.001

The amplitude of the interferon response also varied across doses. The response to the first dose was relatively modest, while the second and third doses elicited more robust responses. Notably, the third dose induced a response of similar magnitude to the second dose, despite the longer interval between doses.

These observations suggest both qualitative and quantitative changes in interferon responses following repeated mRNA vaccination. The similarity in timing between the first and third dose responses, coupled with the increased amplitude of the third dose response, may reflect contributions from both innate and adaptive immunity - potentially involving innate immune training as well as memory CD8+ T cells and antibody-mediated responses. The robust response to the third dose, despite the extended interval since the second dose, suggests that the capacity for strong interferon responses is maintained over time. These findings have implications for understanding the mechanisms of long-term protection conferred by mRNA vaccines and may inform strategies for booster dose timing and vaccine design.

### A systemic inflammation signature, common to various infectious and autoimmune conditions, is observed post-second and third doses

The A35 module aggregate, which we previously associated with systemic inflammatory responses in subjects with autoimmune and inflammatory diseases (16), was investigated across the three vaccine doses (**Figure 5**). This signature provides insights into the inflammatory component of the immune response to mRNA vaccination.

**Figure 5:**
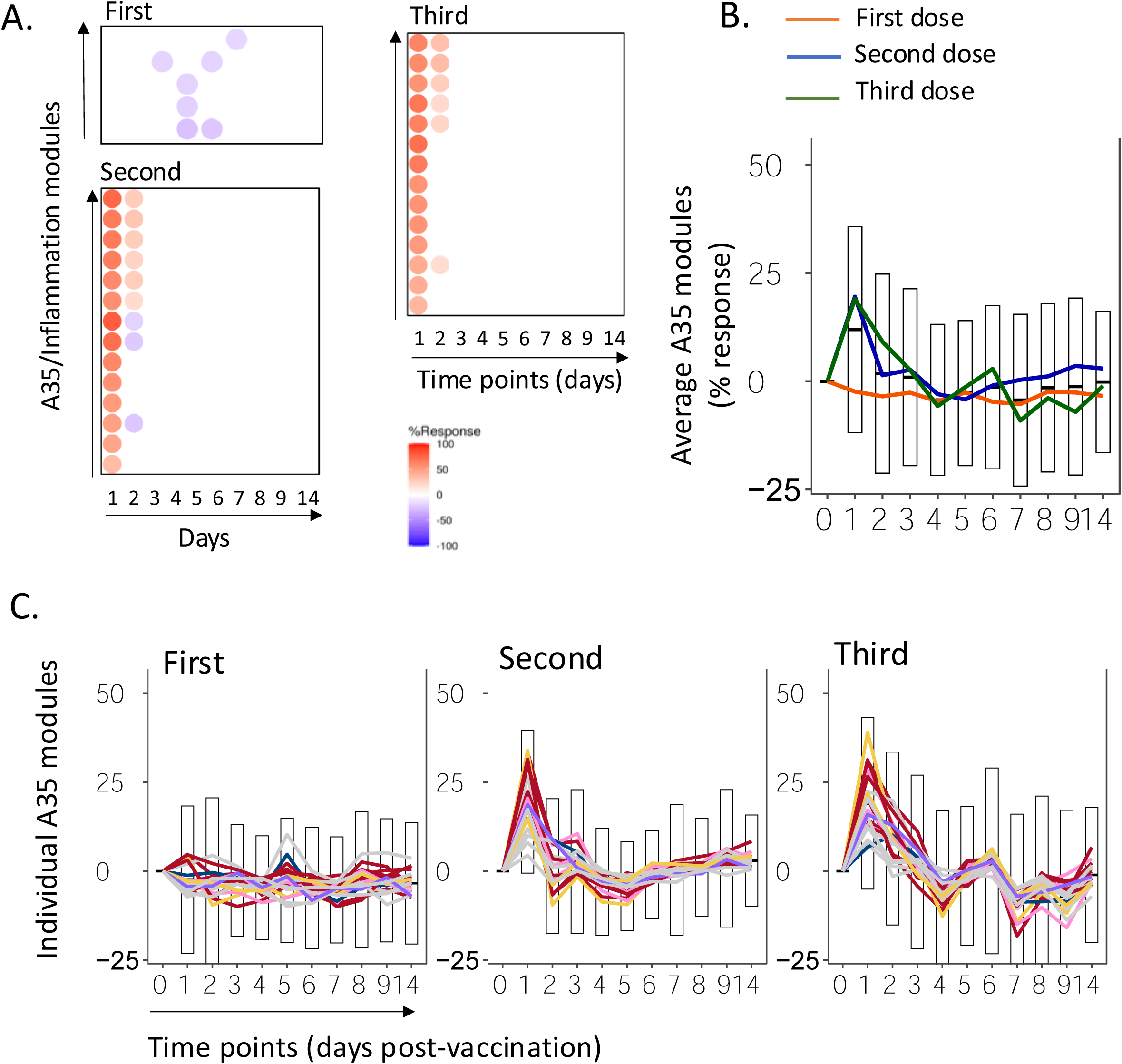
Inflammation response patterns across three doses of COVID-19 mRNA vaccine. (A) Heatmaps showing the response of A35 inflammation modules over time for each vaccine dose. Each row represents one of the A35 modules, and columns represent days post-vaccination. Color intensity indicates the percentage of responsive genes within each module, with red indicating increased expression and blue indicating decreased expression. (B) Line graph showing the average response of all A35 modules over time for each vaccine dose. Lines represent the mean response, with error bars indicating the standard deviation. The orange line represents the first dose, blue the second dose, and green the third dose. (C) Line graphs showing individual A35 module responses for each vaccine dose. Each colored line represents a different module within the A35 aggregate. The gray bars in the background represent the range of responses across all modules.

Following the first dose, the A35 signature did not show an immediate increase (**Figure 5A, 5B**). Instead, a slight decrease was observed after 4-5 days, coinciding with the detection of a subtle adaptive response signature previously reported (12). This pattern suggests a minimal inflammatory response to the initial vaccine exposure. In contrast, the response to the second dose was pronounced, with the A35 signature peaking sharply on day 1 post-vaccination (**Figure 5A, 5B**). This robust inflammatory response indicates a primed innate immune system reacting more vigorously to the second exposure to the vaccine.

The third dose elicited a response of similar amplitude to the second dose, also peaking on day 1 (**Figure 5A, 5B**). However, notable differences in the kinetics of the response were observed. A35 transcript abundance remained significantly elevated on day 2 after the third dose, a time point when post-second dose levels had already returned to baseline. Additionally, a significant decrease in abundance was noticed on days 4 and 7 after the third dose, a trend that was absent following the second dose (**Figure 5B, 5C**). Examination of individual A35 modules revealed consistent patterns across most modules, with some variations in amplitude (**Figure 5C**). This consistency supports the robustness of the aggregate-level observations and suggests a coordinated inflammatory response involving multiple molecular pathways.

These findings indicate that the inflammatory component of the innate immune response to mRNA vaccines is triggered more potently in subjects who have received a priming vaccine dose. The heightened responsiveness persists several months after administration of the second dose, as evidenced by the robust response to the third dose. This pattern mirrors the interferon response signature described earlier, further supporting the concept of innate immune memory or training in the context of mRNA vaccination.

The distinct kinetics of the inflammatory response following the third dose, particularly the prolonged elevation and subsequent decrease, may reflect evolving interactions between innate and adaptive immune components with repeated antigen exposure. These observations contribute to our understanding of how the immune system adapts to repeated mRNA vaccine doses and may have implications for optimizing booster strategies and assessing long-term vaccine efficacy.

### Divergent Temporal Patterns of Erythroid Cell Signature are observed Post-First, Second, and Third Doses

The A37 erythroid cell signature, previously associated with disease severity in RSV infection and elevated in late-stage melanoma patients and liver transplant recipients (17), exhibited distinct patterns across the three vaccine doses (**Figure 6**). Following the first dose, no significant changes were observed in this signature (**Figure 6A, C**). However, the response to the second dose was marked by a sharp peak in transcript abundance on day 1, followed by a decline reaching a maximum decrease on day 5 post-vaccination, before returning to baseline by day 10 (**Figure 6A, B, C**). This day-1 post-second dose increase coincided with the elevation of inflammation-associated modules, representing a novel observation in systems vaccinology studies.

**Figure 6:**
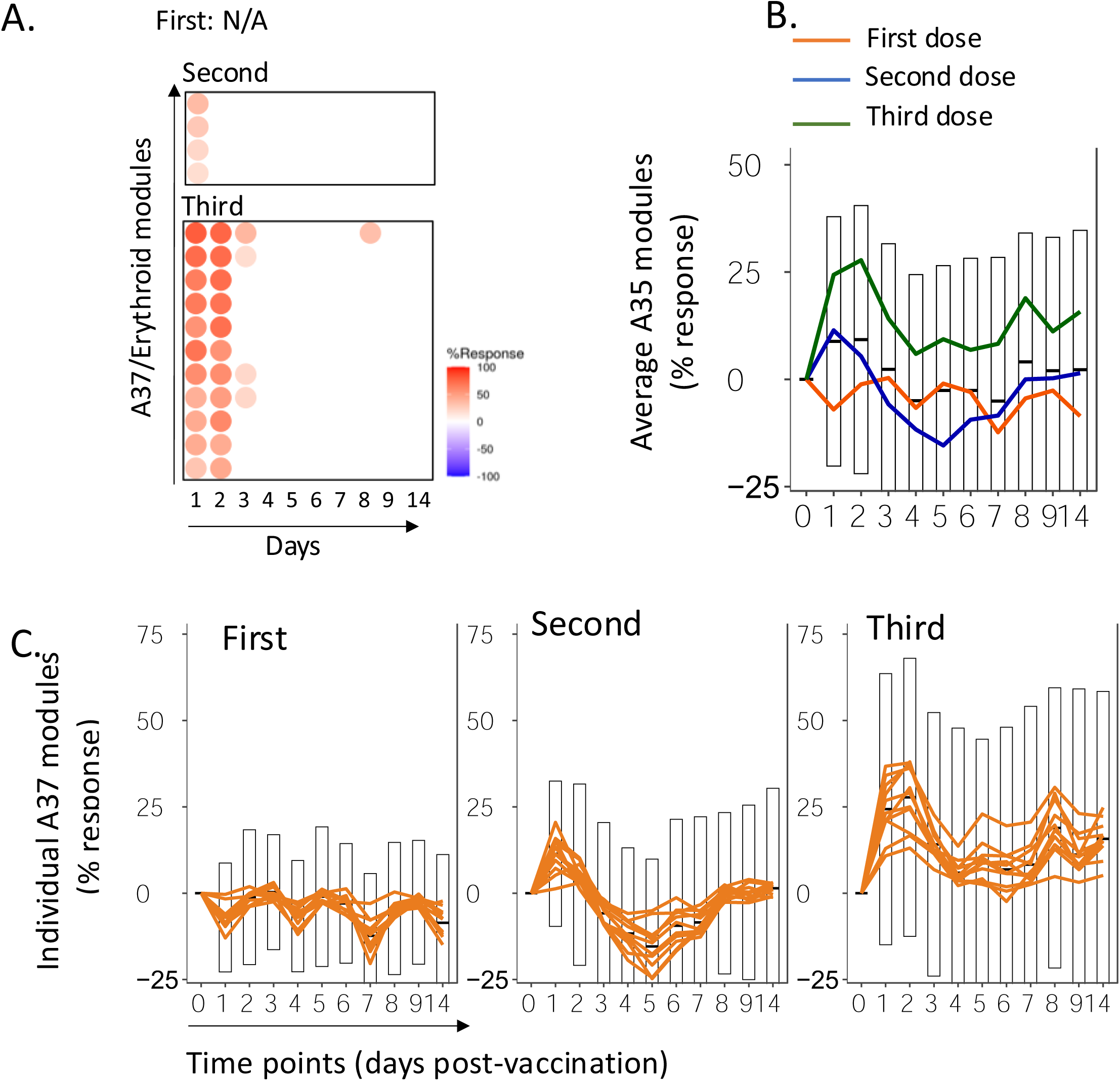
Erythroid cell signature patterns across three doses of COVID-19 mRNA vaccine. (A) Heatmaps showing the response of A37 erythroid modules over time for the second and third vaccine doses. Each row represents one of the A37 modules, and columns represent days post-vaccination. Color intensity indicates the percentage of responsive genes within each module, with red indicating increased expression and blue indicating decreased expression. No significant changes were observed after the first dose (labeled as N/A). (B) Line graph showing the average response of all A37 modules over time for each vaccine dose. The orange line represents the first dose, blue the second dose, and green the third dose. Error bars indicate the standard deviation. (C) Line graphs showing individual A37 module responses for each vaccine dose. Each orange line represents a different module within the A37 aggregate. The gray bars in the background represent the range of responses across all modules. This figure illustrates the divergent temporal patterns of the erythroid cell signature following each dose of the COVID-19 mRNA vaccine, highlighting the contrasting responses observed after the second and third doses, and the unique trajectory of this signature compared to other immune components.

Intriguingly, the response pattern following the third dose differed considerably from that seen after the second dose. While transcript abundance changes were also observed during the first two days post-third dose, the erythroid cell signature trajectory subsequently increased over the week following vaccination (**Figure 6A, B, C**). This trend sharply contrasted with the abundance decrease seen post-second dose over the same period. The divergence in response patterns between the second and third doses was particularly notable, as it differed from the interferon and inflammation signatures, where response kinetics post-third dose largely mirrored those post-second dose.

Erythroid cell populations have been associated with immunosuppressive functions (17–20), suggesting potential implications for the overall immune response to vaccination. The evolving pattern of the erythroid cell signature across vaccine doses highlights the complex and dynamic nature of the immune response to repeated mRNA vaccination. The contrasting trajectories observed after the second and third doses raise intriguing questions about the long-term effects of booster vaccinations on erythropoiesis and potential immunomodulatory consequences. Further investigation into the mechanisms underlying these changes and their potential impact on vaccine efficacy and immune regulation is warranted.

### A robust plasmablast response develops and peaks on day 4 post-third dose

The A27 module aggregate, associated with plasmablast responses, exhibited distinct patterns across the three vaccine doses (**Figure 7**). Following the first dose, a subtle plasmablast signature was detected, as evidenced by the minimal changes in module activity (**Figure 7A, C**). The response amplified considerably after the second dose, with a clear peak observed on day 4 post-vaccination (**Figure 7A, B, C**). This timing is noteworthy, as it precedes the typical day 7 peak observed with other vaccines (21,22), suggesting a potentially accelerated adaptive immune response to mRNA vaccines.

**Figure 7:**
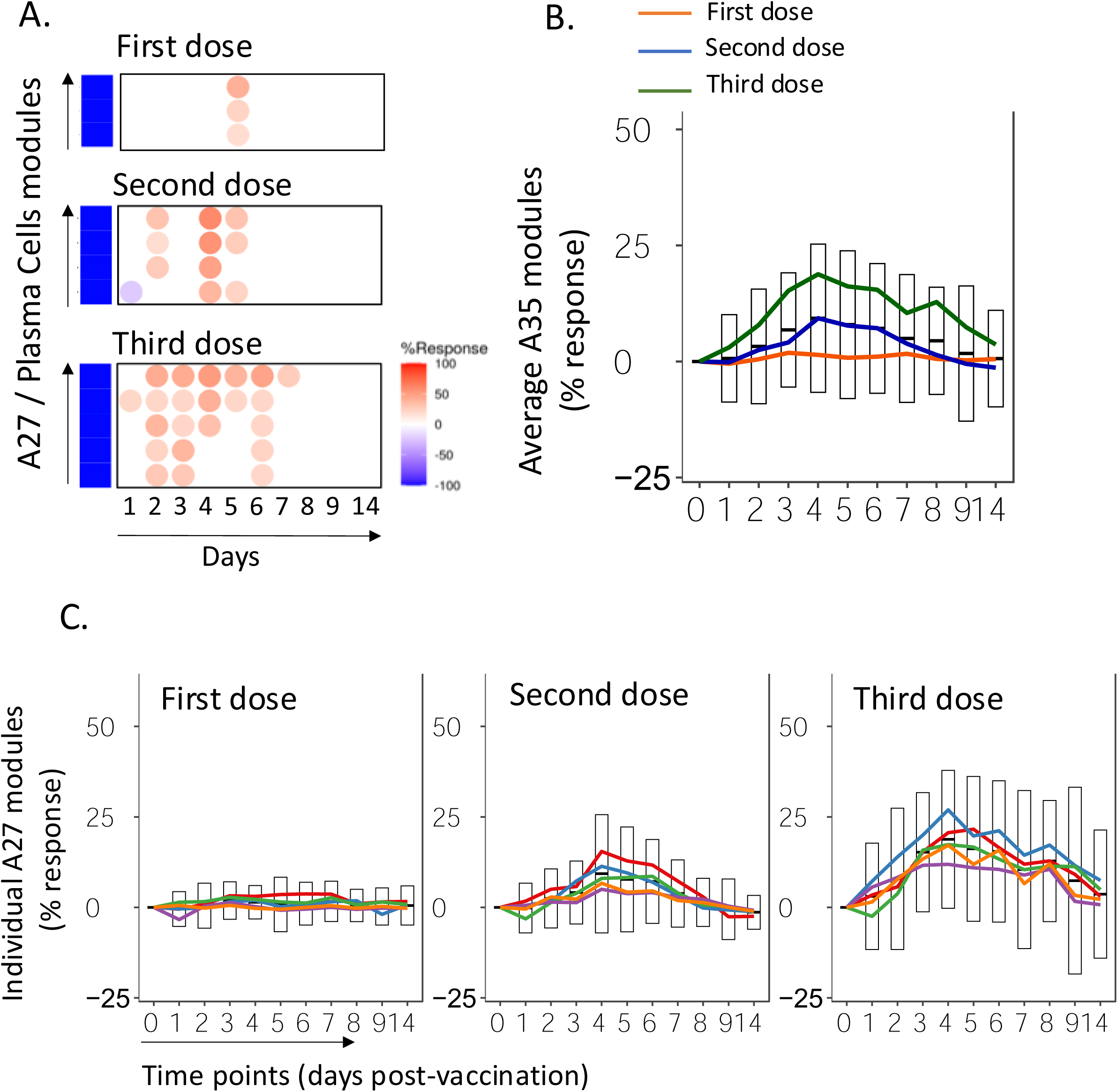
Plasmablast response patterns across three doses of COVID-19 mRNA vaccine. (A) Heatmaps showing the response of A27 plasmablast modules over time for each vaccine dose. Each row represents one of the A27 modules, and columns represent days post-vaccination. Color intensity indicates the percentage of responsive genes within each module, with red indicating increased expression and blue indicating decreased expression. (B) Line graph showing the average response of all A27 modules over time for each vaccine dose. The orange line represents the first dose, blue the second dose, and green the third dose. Error bars (gray) indicate the standard deviation across all modules. (C) Line graphs showing individual A27 module responses for each vaccine dose. Each colored line represents a different module within the A27 aggregate. The gray bars in the background represent the range of responses across all modules.

The plasmablast signature further intensified following the third vaccine dose. The response peaked again on day 4, but with several key differences compared to the second dose response (**Figure 7A, B, C**). First, robust increases in module activity were observed as early as day 2 post-vaccination, indicating a more rapid mobilization of plasmablasts. Second, the magnitude of the peak response was significantly higher, with an average module aggregate response of 21% post-third dose compared to 15% post-second dose [statistical significance to be confirmed]. Finally, the response was more prolonged, returning to baseline only by days 8-9, compared to a faster decline seen after the second dose.

The enhanced plasmablast response following the third dose is particularly significant in light of a recent meta-analysis of blood transcriptome profiles across various vaccines, which identified the peak response of plasmablast signatures as a common determinant of vaccine efficacy (21). The amplified and accelerated response observed here suggests a potent boosting effect of the third vaccine dose on the adaptive humoral response.

This evolving pattern of plasmablast activation across vaccine doses provides valuable insights into the dynamics of B cell responses to repeated mRNA vaccination. The enhanced response to the third dose, despite the extended interval since the second dose, indicates the development of robust immunological memory and suggests that booster doses may be particularly effective in amplifying the adaptive immune response. These findings have important implications for understanding the mechanisms of long-term protection conferred by mRNA vaccines and may inform strategies for optimizing booster dose timing and assessing vaccine efficacy.

## DISCUSSION

In this study, we build upon our previous investigation of immune responses to COVID-19 mRNA vaccines (12). Our earlier work revealed a striking contrast between the responses to the first and second vaccine doses. The first dose elicited a modest response, primarily characterized by an interferon signature, while the second dose induced a robust, polyfunctional response encompassing interferon, inflammatory, and erythroid cell signatures. This dramatic difference in responsiveness raised important questions about the longevity of vaccine-induced immune activation and how the system would respond to subsequent doses administered months later. To address these questions, we employed the same high-resolution temporal profiling approach used in our previous study, analyzing daily blood transcriptome samples for 9 days following administration of a third vaccine dose. This allowed for a direct comparison of immune trajectories across all three doses. Remarkably, we found that the response to the third dose, given on average ∼9 months after the second, was at least as potent as the response observed following the second dose. The third dose elicited strong interferon, inflammatory, and erythroid cell signatures. Interestingly, some aspects of the response kinetics mirrored those seen after the first dose (e.g., the interferon response peaking on day 2), but with amplitudes matching or exceeding those observed after the second dose. Furthermore, we noted an enhanced plasmablast response peaking on day 4, which was further amplified compared to previous doses. These findings provide crucial insights into the evolving nature of immune responses to repeated mRNA vaccination over extended periods. They demonstrate that the heightened responsiveness induced by the initial doses is maintained months later, suggesting effective immune memory and potential long-term efficacy of mRNA vaccine technology. Our results contribute to a more comprehensive understanding of how the immune system adapts to sequential exposures to the same antigen via mRNA vaccination, with implications for optimizing booster strategies and developing next-generation vaccines.

The progressive amplification of the plasmablast response we observed across three doses of mRNA vaccination aligns well with established principles of vaccination and is corroborated by recent studies examining adaptive immune responses to COVID-19 mRNA vaccines. Multiple investigations have demonstrated a stepwise enhancement of both cellular and humoral immunity with each subsequent dose (8,23,24). These studies consistently report increases in neutralizing antibody titers, expansion of memory B cells, and broadening of antibody responses across the three doses. Our transcriptomic evidence of an amplified plasmablast signature mirrors these findings at the molecular level. Furthermore, studies have shown that the third dose not only boosts antibody quantities but also improves antibody quality and breadth (25), supporting our observation of enhanced adaptive responses at the transcriptional level.

Intriguingly, we observed an earlier peak in the plasmablast response (day 4) compared to other vaccines, which may be a unique feature of mRNA vaccines. This finding is supported by Arunachalam et al. (2021), who also noted rapid plasmablast responses following mRNA vaccination (26). The accelerated kinetics of the plasmablast response could be attributed to the direct delivery of mRNA to host cells, bypassing several steps in traditional vaccine platforms. This rapid translation and presentation of antigens may lead to faster activation of B cells and their differentiation into plasmablasts. Moreover, the strong type I interferon response we observed, particularly after the second and third doses, could contribute to this accelerated B cell response, as type I interferons are known to enhance B cell differentiation and antibody production (27). The implications of this earlier plasmablast peak for long-term immunity and vaccine efficacy warrant further investigation, as it may influence the durability and quality of the antibody response.

Our study revealed distinct patterns in the immediate immune response across three doses of mRNA vaccination, particularly in the interferon and inflammatory signatures. These findings both align with and extend current understanding of innate immune activation by mRNA vaccines. The robust interferon response we observed, especially after the second and third doses, is consistent with several recent studies. Arunachalam et al. (2021) reported strong type I interferon signatures within 24 hours of BNT162b2 vaccination, particularly after the second dose (26). Similarly, Kalimuddin et al. (2021) found elevated interferon-stimulated genes following mRNA vaccination (28). Our high-resolution temporal profiling adds nuance to these observations, revealing that the interferon response peaks earlier (day 1) after the second dose compared to the first and third doses (day 2).

The persistence of a strong interferon response even months later with the third dose is particularly noteworthy. This finding suggests a form of innate immune memory or “trained immunity,” a concept gaining traction in vaccine immunology (29). However, the question of antigen specificity remains crucial in determining whether this amplified response can be attributed to adaptive immunity or truly represents trained immunity. Our study design, focused solely on COVID-19 mRNA vaccines, cannot conclusively address this question. As other mRNA vaccines targeting different antigens become approved for human use, comparative studies will be essential to elucidate whether this enhanced responsiveness is antigen-specific (indicating adaptive memory) or a broader phenomenon of innate immune training induced by the mRNA platform itself. Such investigations could have far-reaching implications for our understanding of mRNA vaccine mechanisms and their potential applications beyond COVID-19. The mechanisms underlying this sustained responsiveness warrant further investigation but may involve epigenetic reprogramming of innate immune cells (30), a hallmark of trained immunity.

Our observation of a pronounced inflammatory signature, particularly after the second and third doses, aligns with the reactogenicity profile of mRNA vaccines reported in clinical trials (31). This inflammatory response likely contributes to the vaccines’ immunogenicity but may also explain the increased incidence of side effects after booster doses.

Interestingly, we noted distinct kinetics in the erythroid cell signature across doses, a finding not widely reported in the literature. This signature’s potential role in vaccine-induced immunity or its relation to reported hematological effects of mRNA vaccines (32) merits further exploration.

Our findings underscore the complex and evolving nature of the innate immune response to repeated mRNA vaccination. They highlight the need for further research into the long-term effects of this novel vaccine platform on innate immune function and its implications for vaccine efficacy and safety.

While our study provides valuable insights into the immune responses elicited by three doses of mRNA vaccines, it is important to acknowledge several key limitations. Our relatively small sample size, particularly for the third dose (n=17), may limit the generalizability of our findings. Moreover, we obtained data for all three consecutive doses for only 10 of the original 23 participants from our previous study, potentially introducing bias and limiting our ability to track individual response trajectories comprehensively. Our cohort may not be fully representative of the general population, with factors such as age, sex, ethnicity, pre-existing health conditions, and other potential confounding variables influencing individual responses to vaccination. From a technical perspective, while our high-resolution transcriptomic approach provides a comprehensive view of gene expression changes, it does not directly measure protein levels or cellular functions. The use of whole blood transcriptomics, while broad in scope, may mask cell-type-specific responses. Integration with proteomic, functional immunological assays, and single-cell approaches would provide a more complete picture of the immune response and offer additional resolution to our findings.

In conclusion, our study provides a detailed characterization of the evolving immune response to repeated mRNA vaccination against COVID-19. By employing high-resolution temporal profiling, we have revealed distinct patterns of interferon, inflammatory, erythroid cell, and plasmablast signatures across three vaccine doses. Our findings demonstrate the maintenance and even enhancement of vaccine responsiveness months after the second dose, challenging initial concerns about waning immunity. The observed amplification of both innate and adaptive immune components with each subsequent dose underscores the effectiveness of mRNA vaccines in eliciting robust and increasingly potent immune responses. The unique kinetics of these responses, particularly the accelerated plasmablast peak, may be a distinctive feature of mRNA vaccines and warrants further investigation. While our study has limitations, it provides crucial insights into the complex dynamics of vaccine-induced immunity and lays the groundwork for future, more comprehensive investigations. As mRNA vaccine technology continues to evolve and find applications beyond COVID-19, understanding these immune trajectories will be vital for optimizing vaccination strategies, predicting long-term efficacy, and informing the development of next-generation vaccines. Ultimately, this work contributes to our growing knowledge of how the immune system adapts to repeated antigen exposure via mRNA vaccination, with implications for public health strategies and future vaccine design.

## MATERIALS AND METHODS

### Subject recruitment

Adult subjects eligible for COVID-19 vaccination were enrolled in the study, contingent on their willingness to adhere to the sampling schedule. The study protocol was approved by Sidra Hospital IRB (IRB number 1670047-6), and all participants provided written informed consent. Inclusion criteria aligned with clinical eligibility for vaccination, with the sole exclusion being prior receipt of any COVID-19 vaccine dose.

The cohort for the first and second dose study comprised 23 subjects with a median age of 38 years (range 29-57). Of these, 20 received the Pfizer-BioNTech vaccine and three received the Moderna vaccine.

For the third dose study, we collected high temporal resolution samples from 17 subjects. This included 10 participants from the original cohort and 7 newly recruited subjects. Demographic information, health status at enrollment, and vaccination side effects for all participants in the third dose study are detailed in **Supplementary File 1**. For the first two doses, the average interval was 22.13 days, with a standard deviation of 2.62 days. The interval between the second and third doses was 264.30 days on average, with a standard deviation of 28.58 days.

### Sampling protocol

For transcriptomics applications in the COVAX study, blood was collected using a finger stick method. Briefly, 50 µl of blood was collected in a capillary/microfuge tube assembly supplied by KABE Labortechnik (Numbrecht, Germany) containing 100 µl of tempus RNA-stabilizing solution aliquoted from a regular-sized tempus tube (designed for the collection of 3 ml of blood and containing 6 ml of solution; ThermoFisher, Waltham, MA, USA). This method is described in detail in an earlier report (13), and the collection procedure is illustrated in an uploaded video: https://www.youtube.com/watch?v=xnrXidwg83I.

Blood was collected prior to the vaccine being administered (day 0), on the same day, and daily thereafter over the next 10 days. This protocol was followed for all three vaccine doses.

For serology applications, 20 µl of blood was collected using a Mitra blood collection device (Neoteryx, Torrance, CA, USA) prior to the vaccine being administered and on days 7 and 14 after vaccination with each dose.

### RNA extraction and QC

RNA was extracted from small blood volumes using the Tempus Spin RNA Isolation Kit (ThermoFisher), following previously documented methods (33). DNA contaminants were eliminated using TurboDNAse (ThermoFisher), and RNA quantity and quality were measured using Qubit (ThermoFisher) and Agilent 2100 Bioanalyzer (Agilent, Santa Clara, California, USA).

### RNA sequencing

For the COVAX cohort, mRNA sequencing was performed using the QuantSeq 3’ mRNA-Seq Library Prep Kit FWD for Illumina, generating 75 base pair single-end reads. Sequencing depth was set to 8 million reads per sample, with an average alignment rate of 79.60%. To ensure robust data, each sample was sequenced across four lanes, and the resulting FASTQ files were merged.

Quality control measures included trimming of adapter sequences and polyA tails. Trimmed reads were then aligned to the human genome (GRCh38/hg38, Genome Reference Consortium Human Build 38, INSDC Assembly GCA_000001405.28, Dec 2013) using STAR 2.6.1d. Raw counts were generated using featureCounts v2.0.0. Raw expression data were normalized using the DESeq2 R package to account for size factor effects. Subsequent analyses were conducted using R version 4.1 or later. We assessed global transcriptional differences between samples using principal component analysis with the “prcomp” function.

The transcriptome profiling data, along with detailed sample information, have been deposited in the NCBI Gene Expression Omnibus (GEO) under accession ID GSE281864.

### Statistical Analysis

We conducted analyses using pre-defined gene sets, specifically employing a fixed repertoire of 382 transcriptional modules that were thoroughly functionally annotated, as described in a recent publication (15). This repertoire of transcriptional modules (“BloodGen3”) was identified based on co-expression patterns observed across 16 blood transcriptome datasets encompassing 985 individual transcriptome profiles.

We used a custom R package (34) for downstream analysis and visualization. The workflow consisted of three main steps:

1. Annotating the expression matrix (DESeq2 normalized counts) with module repertoire information, mapping transcripts to BloodGen3 modules.
2. Determining differential expression at either the group or individual sample level.
3. Calculating the “module response,” defined as the percentage of constitutive transcripts with altered abundance between two study groups or for an individual compared to baseline (pre-vaccination levels in this study).

Module response values ranged from 100% (all constitutive transcripts increased) to −100% (all constitutive transcripts decreased). For visualization on fingerprint grids or heatmaps, only the dominant trend (increase or decrease in abundance) was retained, with red indicating an increase and blue a decrease.

For group comparisons, we applied p-value and false discovery rate cutoffs (DESeq2 FDR <0.1). For longitudinal analyses, we used fixed fold-change and expression difference cutoffs (|FC|>1.5 and |DIFF|>10). We determined significance for each module using the differential gene set enrichment function of the dearseq R package (35). * p<0.05, ** p <0.01, *** p < 0.001.

We visualized module responses using fingerprint grid plots and heatmaps, representing responses for groups or individual subjects compared to pre-vaccination baselines. We generated line graphs showing average individual responses over time using the ggplot2 R package (36).

## FUNDING

The work of D.R., S.D., B.S.A.K., G.G., M.T., L.M., L.L., F.R.V., G.M., S.L., T.A., N.V.P., D.B., J.-C.G., and D.C. was supported by Sidra Medicine Internal funds (SDR400162).

### AUTHOR CONTRIBUTIONS

Conceptualization: D.R., S.D., D.B., J.-C.G., and D.C. Methodology: D.R., S.D., B.S.A.K., G.G., D.B., J.-C.G., and D.C. Data generation: D.R., S.D., B.S.A.K., G.G., M.T., L.M., L.L., F.R.V., G.M., S.L., D.B., J.-C.G., and D.C. Investigation: D.R., S.D., D.B., J.-C.G., and D.C. Resources: D.R., S.D., T.A., N.V.P., D.B., J.-C.G., and D.C. Formal analysis: D.R., S.D., B.S.A.K., G.G., M.T., L.M., L.L., F.R.V., G.M., S.L., D.B., J.-C.G., and D.C. Writing—original draft: D.R., S.D., G.G., D.B., J.-C.G., and D.C. Writing—review and editing: all authors. Visualization: D.R., S.D., and D.C. Supervision: D.R., S.D., S.L., D.B., J.-C.G., and D.C.

### DATA AND MATERIALS AVAILABILITY

All data needed to evaluate the conclusions in the paper are present in the paper and/or the Supplementary Materials. RNA-seq data have been made available in the NCBI GEO repository under accession ID GSE281864.

## REFERENCES

1. Baden LR, El Sahly HM, Essink B, Kotloff K, Frey S, Novak R, et al. Efficacy and Safety of the mRNA-1273 SARS-CoV-2 Vaccine. N Engl J Med. 2021 Feb 4;384(5):403–16.

2. Polack FP, Thomas SJ, Kitchin N, Absalon J, Gurtman A, Lockhart S, et al. Safety and Efficacy of the BNT162b2 mRNA Covid-19 Vaccine. N Engl J Med. 2020 Dec 31;383(27):2603–15.

3. Chemaitelly H, Tang P, Hasan MR, AlMukdad S, Yassine HM, Benslimane FM, et al. Waning of BNT162b2 Vaccine Protection against SARS-CoV-2 Infection in Qatar. N Engl J Med. 2021 Dec 9;385(24):e83.

4. Tartof SY, Slezak JM, Fischer H, Hong V, Ackerson BK, Ranasinghe ON, et al. Effectiveness of mRNA BNT162b2 COVID-19 vaccine up to 6 months in a large integrated health system in the USA: a retrospective cohort study. Lancet Lond Engl. 2021 Oct 16;398(10309):1407–16.

5. Dagan N, Barda N, Kepten E, Miron O, Perchik S, Katz MA, et al. BNT162b2 mRNA Covid-19 Vaccine in a Nationwide Mass Vaccination Setting. N Engl J Med. 2021 Apr 15;384(15):1412–23.

6. Watson OJ, Barnsley G, Toor J, Hogan AB, Winskill P, Ghani AC. Global impact of the first year of COVID-19 vaccination: a mathematical modelling study. Lancet Infect Dis. 2022 Sep;22(9):1293–302.

7. Sahin U, Muik A, Derhovanessian E, Vogler I, Kranz LM, Vormehr M, et al. COVID-19 vaccine BNT162b1 elicits human antibody and TH1 T cell responses. Nature. 2020 Oct;586(7830):594–9.

8. Goel RR, Painter MM, Apostolidis SA, Mathew D, Meng W, Rosenfeld AM, et al. mRNA vaccines induce durable immune memory to SARS-CoV-2 and variants of concern. Science. 2021 Dec 3;374(6572):abm0829.

9. Arunachalam PS, Walls AC, Golden N, Atyeo C, Fischinger S, Li C, et al. Adjuvanting a subunit COVID-19 vaccine to induce protective immunity. Nature. 2021 Jun;594(7862):253–8.

10. Stephenson E, Reynolds G, Botting RA, Calero-Nieto FJ, Morgan MD, Tuong ZK, et al. Single-cell multi-omics analysis of the immune response in COVID-19. Nat Med. 2021 May;27(5):904–16.

11. Qi F, Cao Y, Zhang S, Zhang Z. Single-cell analysis of the adaptive immune response to SARS-CoV-2 infection and vaccination. Front Immunol. 2022;13:964976.

12. Rinchai D, Deola S, Zoppoli G, Kabeer BSA, Taleb S, Pavlovski I, et al. High-temporal resolution profiling reveals distinct immune trajectories following the first and second doses of COVID-19 mRNA vaccines. Sci Adv. 2022 Nov 11;8(45):eabp9961.

13. Rinchai D, Anguiano E, Nguyen P, Chaussabel D. Finger stick blood collection for gene expression profiling and storage of tempus blood RNA tubes [Internet]. F1000Research; 2017 [cited 2025 Jan 3]. Available from: https://f1000research.com/articles/5-1385

14. Srivastava K, Carreño JM, Gleason C, Monahan B, Singh G, Abbad A, et al. SARS-CoV-2-infection- and vaccine-induced antibody responses are long lasting with an initial waning phase followed by a stabilization phase. Immunity. 2024 Mar 12;57(3):587–599.e4.

15. Altman MC, Rinchai D, Baldwin N, Toufiq M, Whalen E, Garand M, et al. Development of a fixed module repertoire for the analysis and interpretation of blood transcriptome data. Nat Commun. 2021 Jul 19;12(1):4385.

16. Rawat A, Rinchai D, Toufiq M, Marr AK, Kino T, Garand M, et al. A Neutrophil-Driven Inflammatory Signature Characterizes the Blood Transcriptome Fingerprint of Psoriasis. Front Immunol. 2020;11:587946.

17. Rinchai D, Altman MC, Konza O, Hässler S, Martina F, Toufiq M, et al. Definition of erythroid cell-positive blood transcriptome phenotypes associated with severe respiratory syncytial virus infection. Clin Transl Med. 2020 Dec;10(8):e244.

18. Namdar A, Koleva P, Shahbaz S, Strom S, Gerdts V, Elahi S. CD71+ erythroid suppressor cells impair adaptive immunity against Bordetella pertussis. Sci Rep. 2017 Aug 10;7(1):7728.

19. Grzywa TM, Sosnowska A, Rydzynska Z, Lazniewski M, Plewczynski D, Klicka K, et al. Potent but transient immunosuppression of T-cells is a general feature of CD71+ erythroid cells. Commun Biol. 2021 Dec 10;4(1):1384.

20. Long H, Jia Q, Wang L, Fang W, Wang Z, Jiang T, et al. Tumor-induced erythroid precursor-differentiated myeloid cells mediate immunosuppression and curtail anti-PD-1/PD-L1 treatment efficacy. Cancer Cell. 2022 Jun 13;40(6):674–693.e7.

21. Hagan T, Gerritsen B, Tomalin LE, Fourati S, Mulè MP, Chawla DG, et al. Transcriptional atlas of the human immune response to 13 vaccines reveals a common predictor of vaccine-induced antibody responses. Nat Immunol. 2022 Dec;23(12):1788–98.

22. Obermoser G, Presnell S, Domico K, Xu H, Wang Y, Anguiano E, et al. Systems scale interactive exploration reveals quantitative and qualitative differences in response to influenza and pneumococcal vaccines. Immunity. 2013 Apr 18;38(4):831–44.

23. Muecksch F, Wang Z, Cho A, Gaebler C, Ben Tanfous T, DaSilva J, et al. Increased memory B cell potency and breadth after a SARS-CoV-2 mRNA boost. Nature. 2022 Jul;607(7917):128–34.

24. Röltgen K, Nielsen SCA, Silva O, Younes SF, Zaslavsky M, Costales C, et al. Immune imprinting, breadth of variant recognition, and germinal center response in human SARS-CoV-2 infection and vaccination. Cell. 2022 Mar 17;185(6):1025–1040.e14.

25. Gruell H, Vanshylla K, Tober-Lau P, Hillus D, Schommers P, Lehmann C, et al. mRNA booster immunization elicits potent neutralizing serum activity against the SARS-CoV-2 Omicron variant. Nat Med. 2022 Mar;28(3):477–80.

26. Arunachalam PS, Scott MKD, Hagan T, Li C, Feng Y, Wimmers F, et al. Systems vaccinology of the BNT162b2 mRNA vaccine in humans. Nature. 2021 Aug;596(7872):410–6.

27. Proietti E, Bracci L, Puzelli S, Di Pucchio T, Sestili P, De Vincenzi E, et al. Type I IFN as a natural adjuvant for a protective immune response: lessons from the influenza vaccine model. J Immunol Baltim Md 1950. 2002 Jul 1;169(1):375–83.

28. Kalimuddin S, Tham CYL, Qui M, de Alwis R, Sim JXY, Lim JME, et al. Early T cell and binding antibody responses are associated with COVID-19 RNA vaccine efficacy onset. Med N Y N. 2021 Jun 11;2(6):682–688.e4.

29. Netea MG, Domínguez-Andrés J, Barreiro LB, Chavakis T, Divangahi M, Fuchs E, et al. Defining trained immunity and its role in health and disease. Nat Rev Immunol. 2020 Jun;20(6):375–88.

30. Saeed S, Quintin J, Kerstens HHD, Rao NA, Aghajanirefah A, Matarese F, et al. Epigenetic programming of monocyte-to-macrophage differentiation and trained innate immunity. Science. 2014 Sep 26;345(6204):1251086.

31. Polack FP, Thomas SJ, Kitchin N, Absalon J, Gurtman A, Lockhart S, et al. Safety and Efficacy of the BNT162b2 mRNA Covid-19 Vaccine. N Engl J Med. 2020 Dec 31;383(27):2603–15.

32. Lee EJ, Cines DB, Gernsheimer T, Kessler C, Michel M, Tarantino MD, et al. Thrombocytopenia following Pfizer and Moderna SARS-CoV-2 vaccination. Am J Hematol. 2021 May 1;96(5):534–7.

33. Kabeer BSA, Tomei S, Mattei V, Brummaier T, McGready R, Nosten F, et al. A protocol for extraction of total RNA from finger stick whole blood samples preserved with Tempus^TM^ solution [Internet]. F1000Research; 2018 [cited 2025 Jan 3]. Available from: https://f1000research.com/articles/7-1739

34. Rinchai D, Roelands J, Toufiq M, Hendrickx W, Altman MC, Bedognetti D, et al. BloodGen3Module: blood transcriptional module repertoire analysis and visualization using R. Bioinforma Oxf Engl. 2021 Aug 25;37(16):2382–9.

35. Agniel D, Hejblum BP. Variance component score test for time-course gene set analysis of longitudinal RNA-seq data. Biostat Oxf Engl. 2017 Oct 1;18(4):589–604.

36. Wickham H. Data Analysis. In: Wickham H, editor. ggplot2: Elegant Graphics for Data Analysis [Internet]. Cham: Springer International Publishing; 2016 [cited 2025 Jan 3]. p. 189–201. Available from: 10.1007/978-3-319-24277-4_9

